# Reduced Synaptic Heterogeneity in a Tetanus Toxin Model of Epilepsy: Insights from Computational Modeling

**DOI:** 10.1101/2024.09.30.615990

**Authors:** Parvin Zarei Eskikand, Mark J. Cook, Anthony N. Burkitt, David B. Grayden

## Abstract

A neural mass model was used to assess connectivity strength across diverse populations by fitting the model to background EEG data obtained from a Tetanus Toxin rat model of epilepsy. Our findings reveal a notable decline in the variability of estimated parameters when using EEG data recorded from rats in the Tetanus Toxin group compared with the control group. A detailed comparison of standard deviations in estimated parameters between day 1 and day 20 recordings, coinciding with a heightened number of seizures, underscores the impact of Tetanus Toxin on diminishing synaptic strength variability across recordings. This study supports electrophysiological studies suggesting that epileptogenesis induces a reduction in biophysical heterogeneity, potentially leading to an increase in network synchrony associated with epilepsy. Furthermore, our computational model establishes a foundation for future explorations of the implications of this diminished variability.

## I. Introduction

Epilepsy, a multifaceted neurological disorder characterized by recurrent seizures, remains a formidable challenge in both clinical and research settings. Understanding the varied dynamics underlying epileptic conditions is crucial for advancing therapeutic interventions and enhancing our understanding of the associated neural processes. This study explores the neural dynamics in a Tetanus Toxin (TT) rat model, a unique experimental paradigm that offers insights into the specific context of toxin-induced seizures [5]. Leveraging the power of computational modeling, we employ a neural mass model to investigate connectivity strengths within the cortical column, focusing in this study upon the implications for the stability and uniformity of the estimated parameters under different conditions [7].

We fit the the parameters of a neural mass model to the intracranial EEG recorded from TT and control rats to obtain insights from the estimated connectivity strengths of the model during epileptogenesis. In the subsequent sections, we present our findings, examining the calculated means and standard deviations of the connectivity strength obtained from fitting to the EEG recordings. Using quantitative comparisons between control and epilepsy groups at different time points, we aim to unravel the complex interplay of neural dynamics in response to tetanus toxin-induced seizures.

## II. Materials AND Methods

### A. Neural Mass Model

A Neural Mass Model for a cortical column was used, composed of three interconnected motifs representing layers 2/3, 4, and 5 of the cortex [7]. Each motif consists of an excitatory neural population and an inhibitory neuron population interconnected through forward and recurrent inhibitory and excitatory connections. Excitatory populations receive both excitatory recurrent and inhibitory inputs from within the same motif, while inhibitory populations receive inhibitory recurrent and excitatory inputs from within the same motif.

The neural populations are described using current-based synapses with voltage dynamics given by

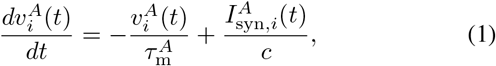

Where 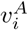 denotes the mean membrane potential of a neuron population, superscript *A* specifies whether the population is excitatory (*E*) or inhibitory (*I*), and subscript *i* represents the layer number (2/3, 4, or 5). The parameter values are listed in Table 1 of [8]. The membrane time constants, 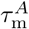, can differ for excitatory and inhibitory neurons, and *c* represents the membrane capacitance. The weighted summation of incoming synaptic currents is given by

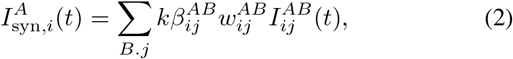

Where 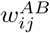 are connectivity weights that are derived from experimental values obtained by the Allen Institute [2, 3, 7], 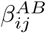 are computed by fitting recorded EEG data to the model output, and *k* is a correction factor aligning values measured in single-cell recordings with the matching operating regime for neural mass models while preserving their relative strengths [7].

Default connectivity weights in the model are derived by considering connection probabilities and synaptic strengths from the Synaptic Physiology database (Allen Institute, USA) [3, 13]. The connectivity weights for intra-columnar connections are detailed in Table 2 of [8].

Post-synaptic current dynamics in the model follow an exponentially decaying function [4, 6],

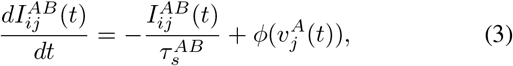

Here, 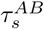 represents the synaptic time constant linked to the specific synapse type between populations *A* and *B*. The values for these synaptic time constants are obtained from recent computational studies, which are fitted to the experimental data [3]. The neural population’s output firing rate is determined by the function *ϕ*(), which involves applying a sigmoid function to the postsynaptic potential,

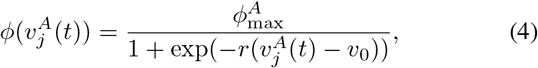

Here, 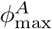 signifies the maximum firing rate, *v*_0_ represents the membrane potential at which the firing rate achieves half of its maximum value, and *r* characterizes the steepness (gain) of the firing rate function.

### B. Fitting the Neural Mass Model to EEG Recordings

The EEG signal output is formulated as the sum of dipole currents (i.e., excitatory to excitatory and inhibitory to excitatory synaptic currents across all cortical layers), multiplied by the membrane resistance. The observational equation takes the form

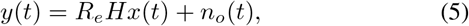

where *y*(*t*) represents the modeled EEG, *R*_*e*_ is a scaling constant, and *H* is the observation vector containing values of 0, 1, and −1, denoting no connection, excitatory, and inhibitory synapses, respectively. The observation noise, *n*_*o*_(*t*) ∼ *N* (0, *σ*), follows a normal distribution with a variance of *σ*.

The objective is to estimate the parameters 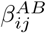 of the model and the states (i.e., membrane potentials and synaptic currents) by fitting the model output to EEG signals recorded from rats in control and TT groups. The 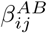 parameters scale default connectivity weights in Table 2 of [8]; is a vector encompassing all 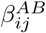 parameters requiring estimation.

Addressing the challenge of multiple solutions generated by optimization algorithms, we introduce parameter dynamics. These dynamics constrain optimization solutions closer to biologically plausible parameter ranges obtained from neurophysiological experiments. The dynamics are expressed as

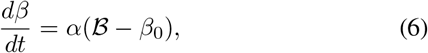

where *β*_0_ is a vector of ones, serving as a fixed-point to guide 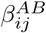 values toward 1 with a decay constant of *α*. This approach ensures that estimated parameters remain proximate to default connectivity weights while minimizing output estimation errors, preventing parameters from drifting over time.

For simultaneous parameter 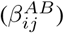 and state estimation, they are combined into a single augmented vector, *x*. The model equation in matrix notation is then given by

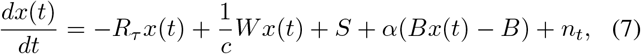

where *x* represents the augmented state vector comprising membrane potentials, synaptic currents, and 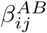 parameters to be estimated. *R*_*τ*_ is a vector containing membrane and synaptic time constants. *W* is a matrix containing default connectivity weights scaled by 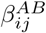 parameters. *S* is a vector of sigmoid function elements associated with state vector dynamics. *B* is a vector indicating the presence of 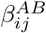 parameters in *x. n*_*t*_ is a vector representing model noise with variance *Q*. Estimation of parameters and states employs the unscented Kalman filter [9]. Details of the applied methods are given in [8].

### C. Data

The data for this investigation were sourced from a Tetanus Toxin model of epilepsy and control rats [5]. In this experiment, six rats constituted the TT group that received intra-hippocampal injections of TT, while the control group comprised four rats that were administered phosphate-buffered saline. To facilitate intracranial EEG recording, each rat was equipped with five stainless steel electrodes implanted into the skull. The onset of spontaneous seizures within the TT group occurred approximately 1-2 weeks post-injection, with the frequency of seizures fluctuating over a period of about 6 weeks. Notably, around 4 to 5 weeks post-injection, there was an observable tendency for the seizure rate to decline until no further seizures were detected. Throughout the experiment, EEG data was continuously recorded at a sampling rate of 2048 Hz for 23 hours each day, with 1 hour designated for routine maintenance checks and data backup. Figure 1 illustrates the changes in the number of seizures over 40 days in a single rat.

**Fig 1.**
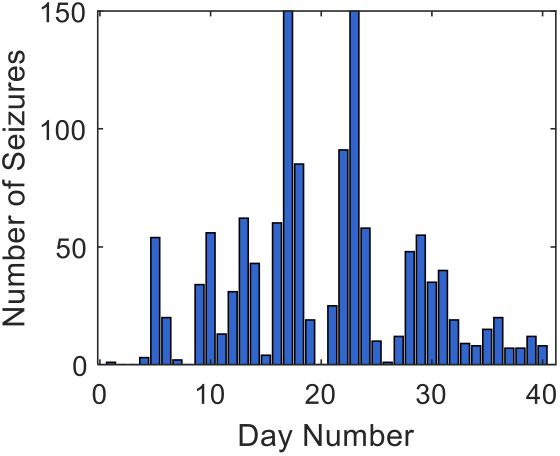
The histogram depicts the changes in the number of seizures from day 1 after the injection of TT to day 40 in one rat.

## III. Results

Fig. 2 displays a sample of the estimated EEG waveform compared with the original EEG recordings. Notably, a close correspondence is observed between the estimated EEG and the original waveform. The unscented Kalman filter demonstrates commendable capability for signal estimation, except in capturing very high-frequency oscillations.

**Fig 2.**
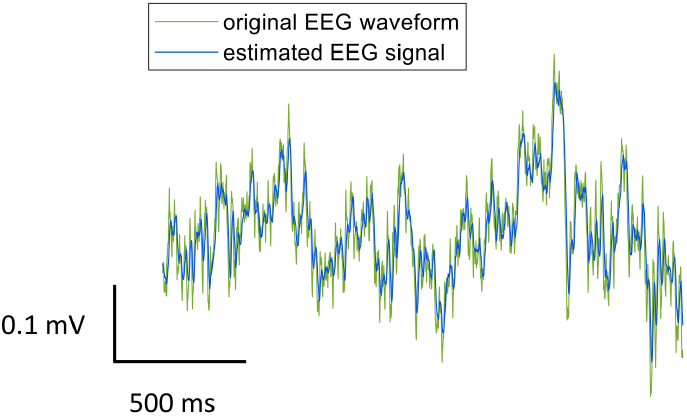
Estimated EEG signal (blue) compared to the original EEG waveform (green) recorded from one rat

We calculated the standard deviations of the connectivity strengths estimated from recordings on day 20 across four rats in the control group and compared them with the standard deviations of parameters obtained from recordings on day 1 and day 20 across four rats in the epilepsy group. For a fair comparison between the standard deviations in the epilepsy group and the control group, we consider only the first four out of six rats in the epilepsy group.

The results of this comparison are shown in Fig. 3. The parameters belonging to each class of control, day 1, and day 2 are connected using dashed lines to make it clear that they belong to one class. The results (Fig. 3) show that the standard deviation of most parameters for the control group were higher than the standard deviations of the parameters on day 20 of the epilepsy group (when the Tetanus Toxin (TT) results in high numbers of seizures). We also compared the parameters on day 20 to those on day 1 for the rats in the epilepsy group. The results show that the standard deviation for day 1 recordings were generally higher than for day 20 recordings for most parameters. The rats in the epilepsy group had far fewer seizures on day 1 of the recordings compared to day 20. We did not find any specific patterns for the mean of the parameters across different groups of rats.

**Fig 3.**
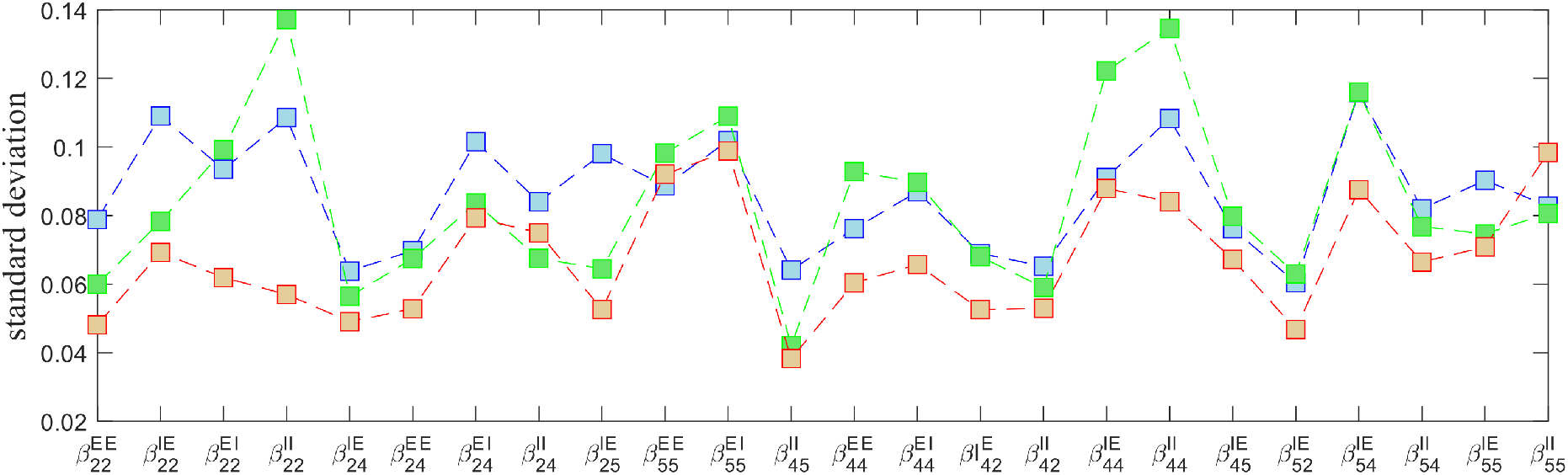
Standard deviation of the model parameters across various recordings and rats. The standard deviation of parameters across different recordings of four control rats (blue) is shown, compared to the standard deviation of parameters across different recordings on day 20 (orange) and day 1 (green) for four rats in the epilepsy group. The x-axis describes different connectivity strengths 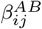 from population *B* in layer *j* to population *A* in layer *i*. The y-axis describes the standard deviation of the parameters across different recordings and rats.

The distribution of standard variation across all parameters and among three different sets— control, day 1 in TT group, and day 20 in TT group is illustrated in the boxplot shown in Fig. 4. The comparison highlights variations in the distribution of standard deviation of the parameters among these three classes. A series of pairwise t-tests were conducted to assess the statistical significance of differences in standard deviation across the three experimental conditions. The null hypothesis posits that, for each pairwise comparison between control, TT on day 1, and TT on day 20, the standard deviation of these parameters belongs to the same distribution. The results revealed a statistically significant difference between the control group and TT (day20), t(46) = 3.8761, p = 0.0003. Additionally, a significant difference was observed between TT (day1) and TT (day20), t(46) = -2.7072, p = 0.0095.

**Fig 4.**
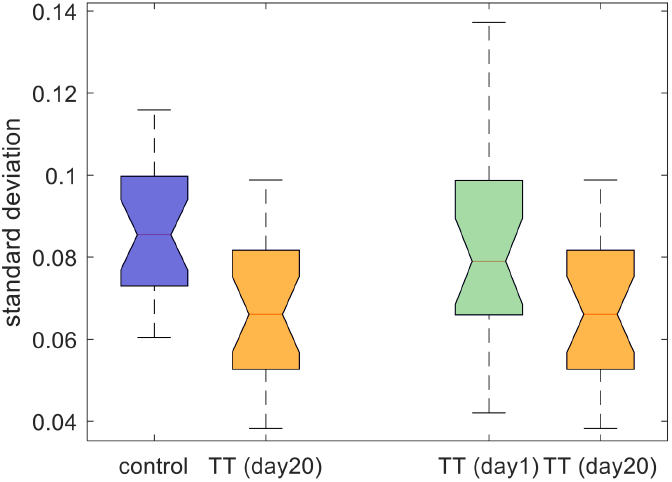
Boxplot of the standard deviation of model parameters across various recordings and rats. Each box represents the InterQuartile Range (IQR) of the respective dataset, with the median indicated by the horizontal line inside the box.

## IV. Discussion AND Conclusion

In this study, a neural mass model of a cortical column was used to fit the connectivity strengths of the model to background (non-seizure) EEG data recorded from Tetanus Toxin and control rats. The optimization results showed a close correspondence between the estimated EEG signals and the original EEG signals. The standard deviation of estimated parameters in the model across different recordings and rats dropped significantly in the case of EEG recorded from TT rats on day 20 compared to control rats (p-value of 0.0003). A higher standard deviation in the control group suggests naturally occurring variability in neural dynamics among healthy rats.

When comparing standard deviations between the control and epilepsy groups on day 20, insights into how the introduction of TT impacts the stability of neural parameters emerge. A lower standard deviation in the epilepsy (TT) group may indicate a more uniform response to toxin-induced seizures. Furthermore, the comparison between standard deviations on day 20 and day 1 offers insights into the temporal evolution of neural dynamics in response to TT. A decrease in standard deviation from day 1 to day 20 might signify a convergence of neural responses as seizures become more frequent, potentially indicating a more severe epileptic condition with synchronized neural responses

Neurophysiological studies using dynamic clamp techniques have also shown reduced variability in the synaptic strengths between interneurons in the case of epilepsy [1, 10, 12]. It was observed that increasing variability in synaptic parameters among given populations of interneurons can significantly reduce the level of synchrony in the network [1, 10, 12]. This effect occurs even in the absence of changes in mean values [12]. Reduced variation appears to be a crucial factor in controlling the excitability of the network and preventing disorders such as epilepsy, which are associated with heightened synchrony in neuronal networks [10].

In a recent study by Rich and colleagues, the results of patch-clamp recordings of layer 5 in epileptogenic and non-epileptogenic tissue demonstrated a significant reduction in biophysical heterogeneity in seizure-generating areas [11]. The results of computational work using spiking networks showed that models characterized by high heterogeneity, mirroring the natural physiological conditions, displayed a more stable asynchronous firing state, which withstands abrupt shifts into a more active and synchronous state (seizure) [11].

These neurophysiological findings and the outcomes of our model support the hypothesis that a pathological reduction in the heterogeneity of synaptic strength across different populations of neurons could result in conditions leading to hypersynchrony in neuronal activity. Our modeling suggests that the difference between the control and epilepsy (TT) groups lies in part upon the standard deviation of parameters in different recordings, not necessarily in the changes in the mean of these parameters. Future research could investigate factors that lead to the loss of heterogeneity in the synaptic properties of the neurons in epilepsy. Understanding the underlying mechanisms and triggers for this reduction in synaptic variability is crucial for unraveling the variability of epileptogenesis and synaptic dysfunction.

## Acknowledgments

The authors thank Dr Warwick Cheung for providing the rat recordings. This work was funded by the Australian Government, under the Australian Research Council’s Training Centre in Cognitive Computing for Medical Technologies.

